# Gabapentin Disrupts Binding of Perlecan to the α_2_δ_1_ Voltage Sensitive Calcium Channel Subunit and Impairs Skeletal Mechanosensation

**DOI:** 10.1101/2022.07.20.500827

**Authors:** Perla C. Reyes Fernandez, Christian S. Wright, Adrianna N. Masterson, Xin Yi, Tristen V. Tellman, Andrei Bonteanu, Katie Rust, Megan L. Noonan, Kenneth E. White, Karl J. Lewis, Uma Sankar, Julia M. Hum, Gregory Bix, Danielle Wu, Alexander G. Robling, Rajesh Sardar, Mary C. Farach-Carson, William R. Thompson

**Author notes:** Corresponding Author: William R. Thompson, DPT, PhD. **Disclosures:** KEW receives royalties for licensing FGF23 to Kyowa Hakko Kirin Co., Ltd; had previous funding from Akebia, and current funding from Calico Labs. KEW also owns equity interest in FGF Therapeutics. All other authors have nothing to disclose.

## Abstract

Our understanding of how osteocytes, the principal mechanosensors within bone, sense and perceive force remains unclear. Previous work identified “tethering elements” (TEs) spanning the pericellular space of osteocytes and transmitting mechanical information into biochemical signals. While we identified the heparan sulfate proteoglycan perlecan (PLN) as a component of these TEs, PLN must attach to the cell surface to induce biochemical responses. As voltage-sensitive calcium channels (VSCCs) are critical for bone mechanotransduction, we hypothesized that PLN binds the extracellular α_2_δ_1_ subunit of VSCCs to couple the bone matrix to the osteocyte membrane. Here, we showed co-localization of PLN and α_2_δ_1_ along osteocyte dendritic processes. Additionally, we quantified the molecular interactions between α_2_δ_1_ and PLN domains and demonstrated for the first time that α_2_δ_1_ strongly associates with PLN via its domain III. Furthermore, α_2_δ_1_ is the binding site for the commonly used pain drug, gabapentin (GBP), which is associated with adverse skeletal effects when used chronically. We found that GBP disrupts PLN::α_2_δ_1_ binding *in vitro*, and GBP treatment *in vivo* results in impaired bone mechanosensation. Our work identified a novel mechanosensory complex within osteocytes composed of PLN and α_2_δ_1_, necessary for bone force transmission and sensitive to the drug GBP. This work provides insights into the mechanisms underlying mechanotransduction and will inform future studies to understand the mechanisms responsible for the negative effects of GBP on bone.

## Introduction

Osteocytes reside deep within the mineralized matrix of bone and have long dendrite-like processes that run through microscopic channels called canaliculi^1^. As osteocytes are uniquely positioned in the bone matrix to communicate with other bone cell types via paracrine signaling and through direct contact with the cellular processes, they are considered the primary mechanosensory skeletal cells^1, 2^. Transmission of mechanical force from the bone matrix to the osteocyte cell membrane was initially thought to occur via direct sensing of whole-tissue strains on the osteocyte surface. However, strains applied to whole bone *in vivo* during normal locomotion are typically between 0.04-0.3% (ref. 3,4), an order of magnitude smaller than the strain necessary to elicit a biochemical response at the osteocyte plasma membrane (1-10%) (ref. 5-7). Thus, a mechanism other than direct force transmission from the bone matrix must account for the ability of osteocytes to perceive mechanical input.

The pericellular space between (PCS) the bone matrix and the osteocyte plasma membrane contains non-mineralized extracellular matrix molecules, including proteoglycans, which are collectively termed the pericellular matrix (PCM)^8, 9^. To explain the mechanism by which tissue-level mechanical strains are transmitted into biochemical responses in osteocytes, the presence of matrix-based "tethering elements" (TEs) able to span the PCS and anchor the osteocyte processes to the mineralized matrix was proposed^10^. This theoretical model was followed by ultrastructural studies using electron microscopy that visually revealed the tethering elements within the PCS^11^. However, the molecular identity of these TEs remained unknown.

Using immunostaining and immunogold assays, we showed that the large heparan sulfate proteoglycan perlecan (HSPG2, PLN) is expressed along osteocyte cell bodies and dendritic processes in cortical bone but not within the mineralized matrix^12^. Furthermore, PLN-deficient mice had fewer TEs within osteocyte canaliculi^12^, lower canalicular drag forces, and decreased responses to exogenous loading^13^. Together, these studies identified PLN as a component of the tethering complex in osteocytes necessary for anabolic responses to mechanical loading. While this finding helped explain how force is transmitted to the osteocyte cell membrane, the PLN-containing tethers must attach to the cell surface to induce biochemical responses.

Intracellular calcium (Ca^2+^) influx is a potent signal in response to force^14^. Ca^2+^ influx is regulated by voltage sensitive Ca^2+^ channels (VSCCs), and *in vitro* and *in vivo* studies have shown that VSCCs are necessary for anabolic responses to skeletal loading^15, 16^. As PLN deficiency impairs mechanically-induced Ca^2+^ signaling in bone^17^, we hypothesized that PLN tethers bind VSCC ectodomains, forming what we call a matrix- channel tethering complex (M-CTC), and that this interaction facilitates intracellular Ca^2+^ influx in response to mechanical force.

VSCCs are integral membrane proteins composed of the pore-forming α_1_ subunit, which enables calcium (Ca^2+^) entry, and auxiliary subunits including α_2_δ, β, and γ^18^ **(Fig. 1)**. While the pore-forming (α_1_) subunit enables Ca^2+^ entry across the membrane, auxiliary subunits influence gating kinetics of the channel pore. In particular, the α_2_δ_1_ subunit is anchored in the plasma membrane, with the majority of the protein positioned extracellularly, an optimal location to interact with extracellular molecules, such as PLN-containing tethering elements. Interestingly, the α_2_δ_1_ subunit is the binding site of the antiepileptic and neuropathic pain drug gabapentin (GBP)^19, 20^ (**Fig. 1**). Chronic GBP use is associated with increased fracture risk in humans^21^ and impaired bone formation in both human and animal studies^22, 23^. However, the mechanism(s) mediating GBP- associated adverse skeletal effects are unclear. Thus, in addition to establishing if PLN directly binds the α_2_δ_1_ subunit of VSCCs, we sought to determine if GBP interferes with binding of the PLN/α_2_δ_1_ complex.

**Figure 1.**
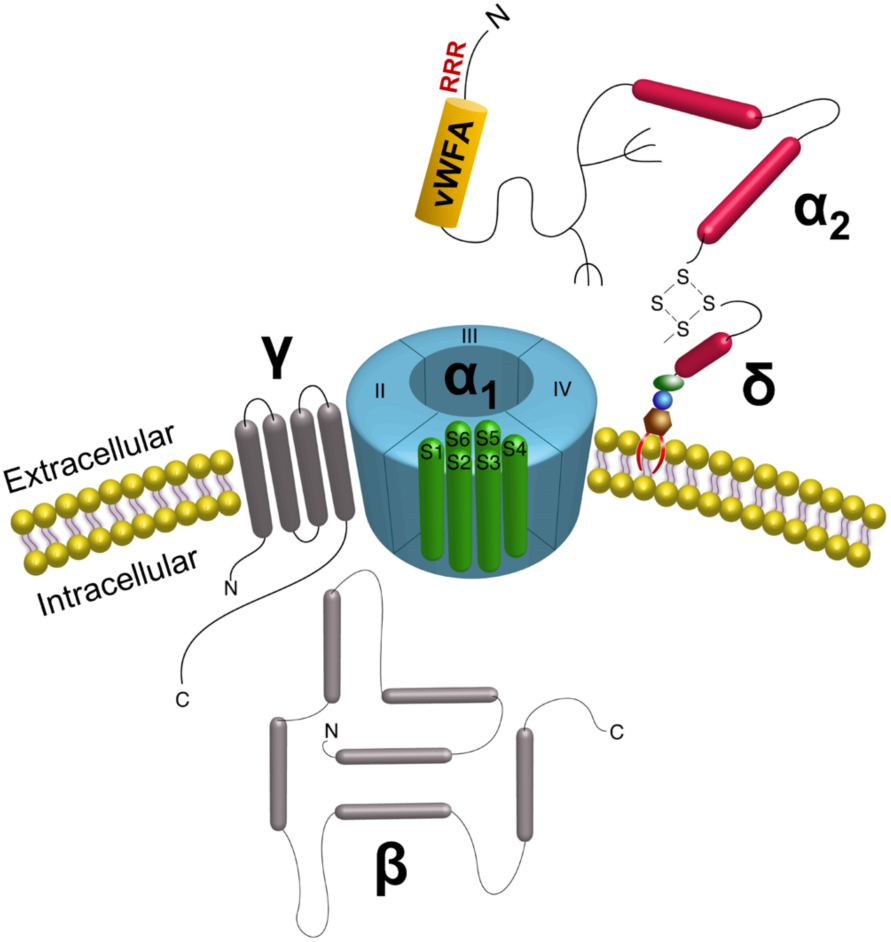
Structure of voltage sensitive calcium channels. The channel complex is composed of the α_1_ pore-forming subunit with auxiliary β, γ, and α_2_δ subunits bound to the pore, positioned to alter gating kinetics of the channel. The α_2_δ subunit is anchored in the membrane via the δ portion, with the α_2_ region positioned extracellularly. In the extracellular portion (α_2_) of the α_2_δ subunit, the von Willebrand Factor A domain (vWFA) sequence and the Arg-Arg-Arg (RRR) motif for Gabapentin binding are indicated. Adapted from Wright *et al.*^81^.

### α_2_δ_1_ and PLN co-localize in murine osteocyte-like cells

We conducted double immunostaining to test whether PLN co-localizes with the pore-forming Ca_v_3.2 (α_1H_) VSCC subunit, with wheat germ agglutinin (WGA), and/or α_2_δ_1_ in MLO-Y4 osteocytic cells. As we previously reported, Ca_v_3.2 (α_1H_) is the primary α_1_ VSCC subunit in osteocytes^24^. In MLO-Y4 cells, Ca_v_3.2 (α_1H_) is expressed within the cell, but also along the cell periphery (**Suppl. Fig. S1a, d**). As WGA binds N-acetyl glucosamine sugars, which are present on the extracellular α_2_ portion of α_2_δ_1_, we performed double staining with Ca_v_3.2 and WGA-FITC (**Suppl. Fig. S1a, b**). Areas of overlap (yellow) validated our previous findings that Ca_v_3.2 associates with α_2_δ_1_ in osteocytes (**Suppl. Fig. S1c**). To determine if PLN associates with Ca_v_3.2 (α_1H_) channels, double staining with Ca_v_3.2 (α_1H_) and PLN was performed (**Suppl. Fig. S1d-f**). Several areas of overlapping signal demonstrated that PLN associates with Ca_v_3.2 (α_1H_) (**Suppl. Fig. S1f**).

Consistent with these findings, PLN and WGA staining overlapped in areas along the cell membrane (**Fig. 2a-c**). Immunostaining of MLO-Y4 cells using antibodies specific to α_2_δ_1_ and PLN, demonstrated that both α_2_δ_1_ (**Fig. 2d**) and PLN (**Fig. 2e**) are produced in osteocytic cells. Merged images showed strong overlapping fluorescent signal of these two proteins (**Fig. 2f**, yellow areas). Importantly, the signal was most prominent along osteocytic processes, demonstrating co-localization of PLN and α_2_δ_1_ in the area of greatest mechanosensitivity (**Fig. 2d-f**). All cell culture immunostaining assays showed no signal when probed with normal IgG in place of the primary antibodies or when using N, N’, N’’-triacetylchitotriose as a negative control for WGA-FITC staining. Consistent with the immunostaining results, co-immunoprecipitation assays using MLO-Y4 lysates, showed that α_2_δ_1_ and PLN interact forming a complex *in vitro* (**Fig. 2g**). Overall, these data suggest that this matrix (PLN)-channel (α_2_δ_1_) tethering complex is a critical component for mechanosensory responses in osteocytes.

**Figure 2.**
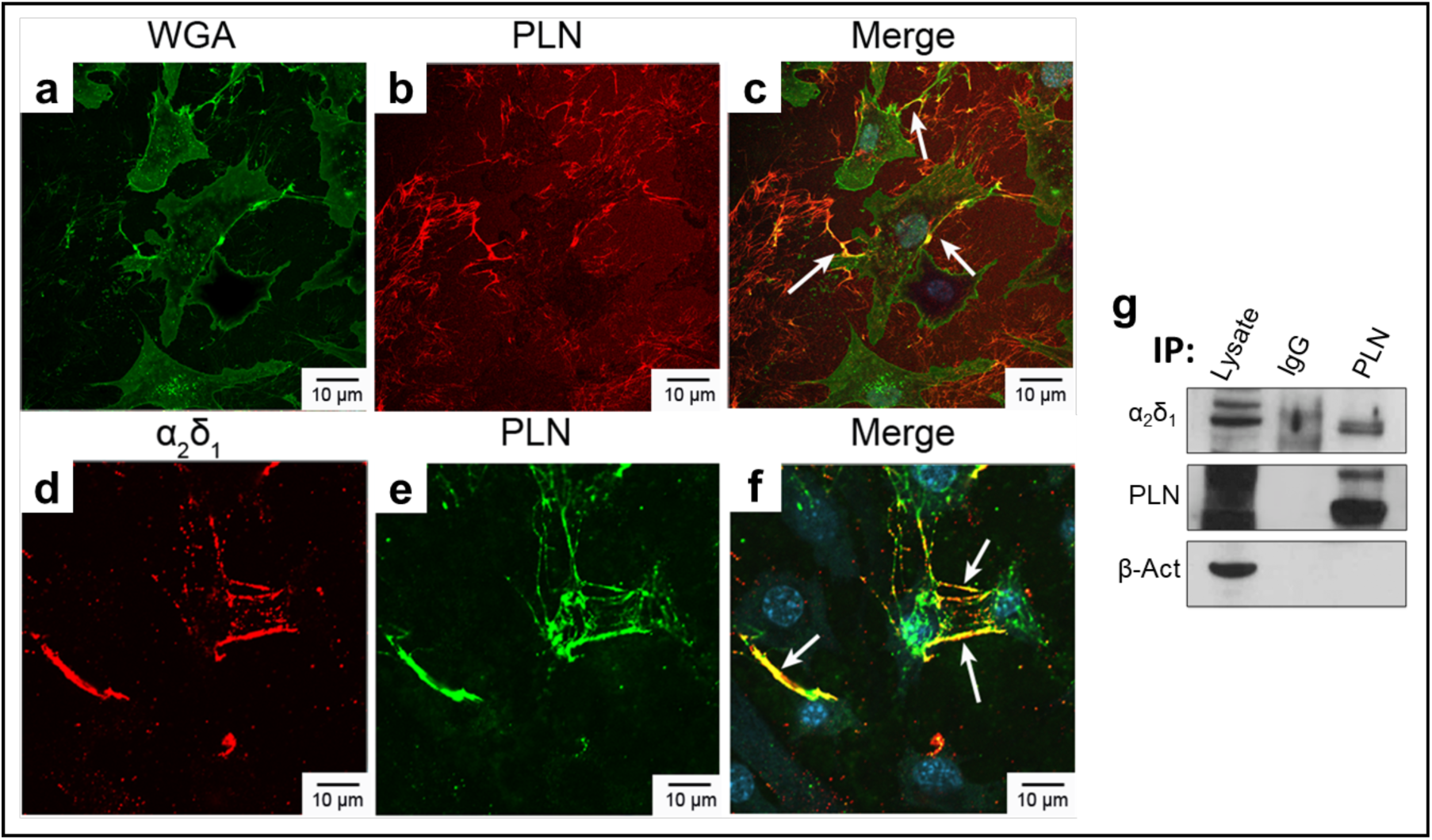
PLN colocalizes with WGA and α_2_δ_1_ in osteocyte-like cells. MLO-Y4 cells stained with **(a)** wheat germ agglutinin (WGA)-FITC (green) and **(b)** perlecan (PLN) (red), **(c)** merge PLN and WGA. On the bottom panels, cells were stained for **(d)** α_2_δ_1_ (red) and **(e)** PLN (green), **(f)** merge PLN and α_2_δ_1_. White arrows in merged images indicate overlapping fluorescent signal. **(g)** Co-immunoprecipitation assays from MLO-Y4 lysates show that PLN and α_2_δ_1_ associate. IgG was used as a negative control. Blots were probed for β-actin antibody as a loading control.

### α_2_δ_1_ and PLN bind with high affinity, which is mediated by PLN Dm III-2

To quantify the molecular interactions between PLN and α_2_δ_1_ we first tested the binding affinity of full-length PLN protein (native form/undigested and enzymatically digested) and the α_2_ portion of α_2_δ_1_, followed by quantifying the binding affinity of individual PLN domains/subdomains (Dm I, III-2, IV-1, IV-2, IV-3, and V) with α_2_. Using LSPR-based experiments (**Suppl. Fig. S2, S3**), we obtained the dissociation constant (the constant describing the drug/receptor interactions at equilibrium) between α_2_-bound sensors and PLN. With a dissociation constant (*K_D_*) of 6.6 x 10^-11^ M, full-length PLN (undigested) bound with high affinity to the α_2_ portion of α_2_δ_1_ (**Table 1**). Removal of heparan sulfate and chondroitin sulfate groups from PLN (digested) resulted in a *K_D_* of 2.6 x 10^-7^ M.

**Table 1.** Binding affinity experiments between the α_2_ portion of the α_2_δ_1_ and perlecan

| Perlecan Domain / Subdomain | $K_D$ (M) |
| --- | --- |
| Undigested Full Length | $6.6 \times 10^{-11}$ |
| Digested Full Length | $2.6 \times 10^{-7}$ |
| Domain I | $7.7 \times 10^{-6}$ |
| **Domain III-2** | **$8.0 \times 10^{-11}$** |
| Domain III-2 (w/cysteine) | $7.7 \times 10^{-6}$ |
| Domain IV-I | $1.4 \times 10^{-7}$ |
| Domain IV-2 | $4.3 \times 10^{-4}$ |
| Domain IV-3 | $1.6 \times 10^{-3}$ |
| Domain V | $5.1 \times 10^{-3}$ |

When examining the individual domains/subdomains of PLN, Dm III-2 had the greatest affinity to the α_2_ polypeptide, displaying a *K_D_* of 8.0 x 10^-11^ M. We also tested binding of α_2_ to Dm III-2 followed by a cysteine-rich sequence. We found that the *K_D_* value for this domain was 7.7 x 10^-6^, suggesting that binding of PLN to α_2_δ_1_ via Dm III-2 is less likely to be mediated through cysteine rich regions. The *K_D_* values of other PLN domains were, Dm I: 7.7 x10^-6^, Dm IV-1: 1.4 x 10^-7^, Dm IV-2: 4.3 x 10^-4^, Dm IV-3: 1.6 x 10^-3^, and Dm V: 5.1 x 10^-3^. These values each demonstrated moderate to weak binding to α_2_ (**Table 1**). The raw data used to obtain the final dissociation constant values are provided as supplementary information (**Suppl. Fig. S4, Suppl. Table S1**).

To evaluate binding of PLN and α_2_δ_1_ *in silico*, computational 3D protein-protein docking models between the von Willebrand Factor A (vWFA) domain of α_2_δ_1_ (4FX5) and PLN domain III-2 (4YEP) were generated. The quality report for structure accuracy confirmed that the models used for receptor (4FX5, 95.7 %) and ligand (4YEP, 100%) had high sequence identities with the input structures, where a sequence ID > 30% is considered reliable. Quality criteria of input protein structures were analyzed by ProQ (v1) (ref. 25), a neural network-based method that predicts the quality of a protein model, as measured by LGscore or MaxSub^26^. Suitable scores for these parameters are classified as correct (LGscore >1.5; MaxSub > 0.1), good (LGscore ≥3 to <5; MaxSub ≥0.5 to <8) or very good (LGscore ≥ 5; MaxSub ≥ 0.8). Input models for receptor 4FX5 (LGscore = 5.77; MaxSub =0.428) and ligand 4YEP (LGscore = 5.811; MaxSub =0.234) were within the appropriate quality ranges for docking modeling. In the HDOCK server, putative binding modes are ranked according to their binding energy scores^27, 28^. The first of the top ten prediction models for 4FX5 and 4YEP scored a docking energy of -272.35, indicating strong protein-protein interactions. Cartoon and surface 3D representations of the highest ranked surface binding prediction model are shown in **Figure 3**.

**Figure 3.**
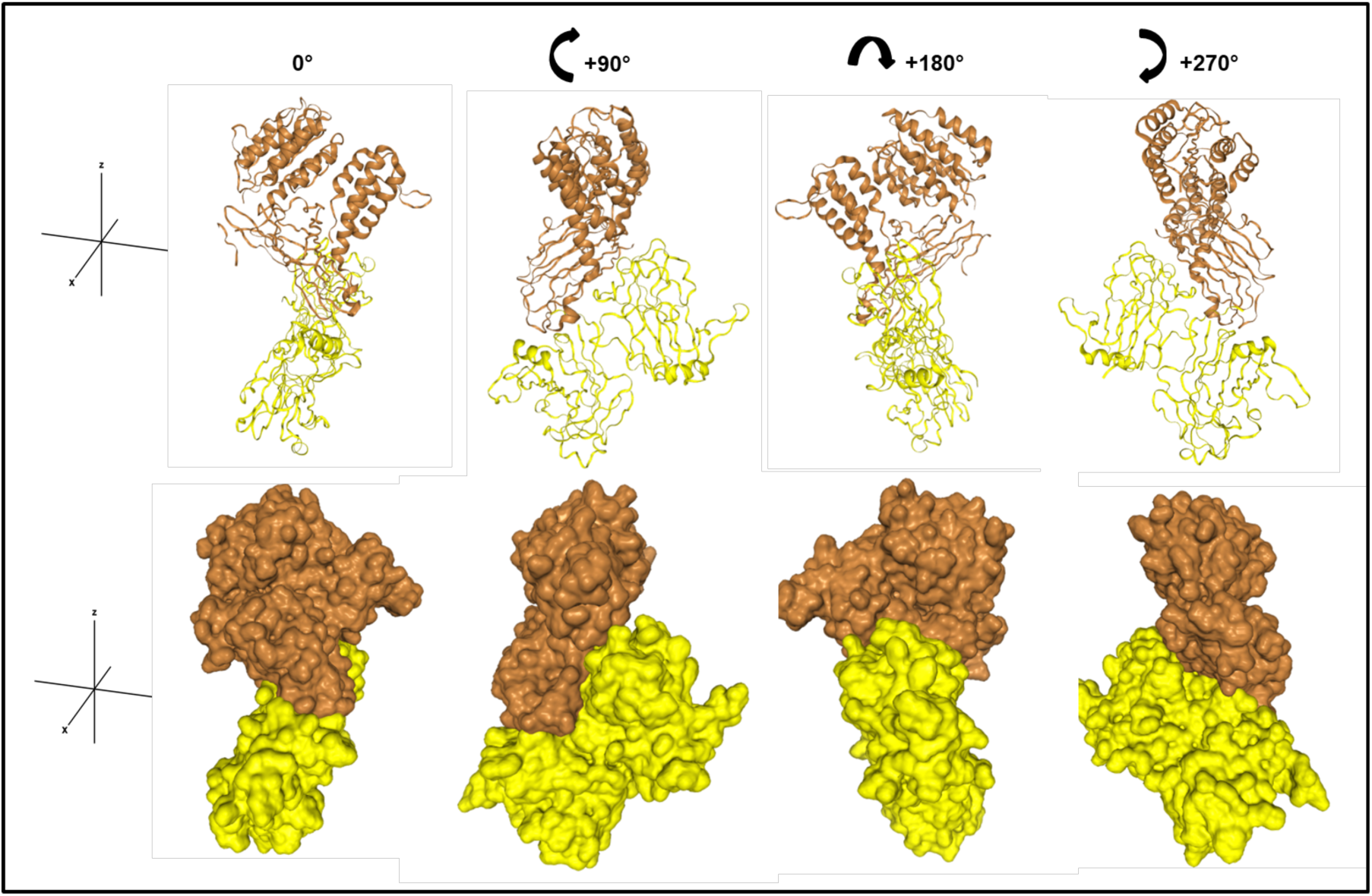
Docking models of vWFA domain of α_2_δ_1_ and PLN Dm III-2. *in silico* protein-protein functional interactions and 3D docking models between the von Willebrand Factor A (vWFA) domain of α_2_δ_1_ and perlecan (PLN) domain (Dm) III-2 were generated with the free web server HDOCK. 4FX5 (brown) is the vWFA domain and 4YEP (yellow) is the L4b domain of human Laminin α_2_ (PLN Dm III-2). Top, cartoon ribbon-style 3D representations of receptor and ligand. Bottom, surface style representation of the proteins. Each image is rotated 90° clockwise from the previous one.

### Gabapentin interferes with PLN:α_2_δ_1_ binding

Since PLN Dm III-2 showed the highest affinity for α_2,_ we then used LSPR assays to determine if binding of PLN Dm III-2 or full-length PLN with α_2_δ_1_ is disrupted by GBP. This was achieved with a series of assays adding either PLN or GBP to the α_2_ peptide bound to the nanoplasmonic sensor. With α_2_ bound to the nanoplasmonic sensor, we first added full-length PLN, which generated a +14.4 nm shift (*Δλ*). Subsequent addition of GBP resulted in a -4.1 nm shift, suggesting dissociation of PLN from α_2_ in the presence of GBP (**Table 2. Exp. 1**).

**Table 2.** LSPR-based interactions among α_2_-functionalized sensors, PLN and GBP

| Exp | Sensors | $\Delta\lambda_{\text{LSPR}}$ (nm) | Added first to the sensors | $\Delta\lambda_{\text{LSPR}}$ (nm) | Added second to the sensors | $\Delta\lambda_{\text{LSPR}}$ (nm) |
| --- | --- | --- | --- | --- | --- | --- |
| 1 | $\alpha_2$ | +39 nm | Full length PLN | +14.4 nm | GBP | -4.1 nm |
| 2 | $\alpha_2$ | +39 nm | GBP | +5.8 nm | Full length PLN | +/- 0.1 nm |
| 3 | $\alpha_2$ | +39 nm | Full length PLN + GBP | +2.7 nm | - | - |
| 4 | $\alpha_2$ | +39 nm | PLN Dm III-2 | +12.7 nm | GBP | -4.3 nm |
| 5 | $\alpha_2$ | +39 nm | GBP | +5.4 nm | PLN Dm III-2 | +0.4 nm |
| 6 | $\alpha_2$ | +39 nm | PLN Dm III-2 + GBP | +4.9 nm | - | - |

Exp = Experiment, LSPR = Localized surface plasmon resonance, PLN = Perlecan, GBP = Gabapentin.

Next, instead of first adding PLN to the nanoprism-bound α_2_ polypeptide, GBP was added which resulted in a +5.8 nm shift. Binding of GBP, then was followed by addition of full-length PLN which generated a shift of only +0.1 nm, indicating an inability of PLN to bind α_2_ in the presence of GBP (**Table 2. Exp. 2**).

Using a third approach, full-length PLN was pre-incubated with GBP, and this combination then was added to the α_2_-bound nanoprism. Addition of the PLN/GBP mixture resulted in a +2.7 nm *Δλ* shift. As the +2.7 nm shift was similar to the +5.8 nm shift observed with GBP binding α_2_ than the +14.4 nm shift found when PLN bound alone, this indicated that in the presence of both GBP and PLN, with equal opportunity to bind, GBP but not PLN bound to the α_2_ polypeptide (**Table 2, Exp. 3**).

A similar series of experiments were conducted to quantify the interactions between PLN Dm III-2, α_2_, and GBP. Here, α_2_ bound with high affinity to Dm III-2 (+12.7 nm shift), and the addition of GBP interfered with this association (-4.3 nm shift) (**Table 2, Exp. 4**). When GBP was bound to α_2_ prior to addition of PLN Dm III-2, the presence of GBP restricted binding of Dm III-2 (+0.4 nm shift) to α_2_ (**Table 2, Exp. 5**), and incubation of α_2_ with a mixture of Dm III-2 and GBP resulted in a shift in the wavelength of +4.9 nm, indicating that GBP bound to α_2_, but not PLN Dm III-2 (**Table 2, Exp. 6**).

### Gabapentin impairs bone mechanosensation *in vivo*

To determine the effects of GBP on skeletal mechanosensitivity we examined changes in anabolic bone responses to mechanical loading in mice treated with GBP or saline (vehicle, VEH). At the time of experiment mice in the VEH and GBP groups had body weights of 29.3 ± 0.41and 29.9 ± 0.43 g, respectively (mean ± SEM). The body weight was not different between groups (p=0.41) and remained stable over the 4 weeks of treatment. In VEH treated mice, as expected, dynamic histomorphometry analyses of loaded ulnas revealed a significant increase in periosteal mineralizing surface (MS/BS) (+17.1 %, p=0.005), bone formation rate (BFR/BS) (+40.4%, p=0.004) and mineral apposition rate (+23.5%, p=0.038) compared to non-loaded controls (**Fig. 4a-b, Table 3)**. In contrast, GBP treatment resulted in blunted bone mechanosensitivity and impaired bone formation. While mice treated with GBP had increased MAR (+17.9% vs non-loaded, p=0.012) there was no change in MS/BS (p=0.67) or BFR/BS (p=0.38) following mechanical loading (**Fig. 4a-b, Table 3)**. The final number of animals included in the analysis was n=9 for the VEH-treated mice and n=7 for the GBP treated mice. One animal from the GBP group was euthanized before completion of the experiment (broken ulna during initial loading) and another mouse from the same group was removed due poor histological quality of control (non-loaded) sections, and thus inability to conduct proper paired comparisons.

**Figure 4.**
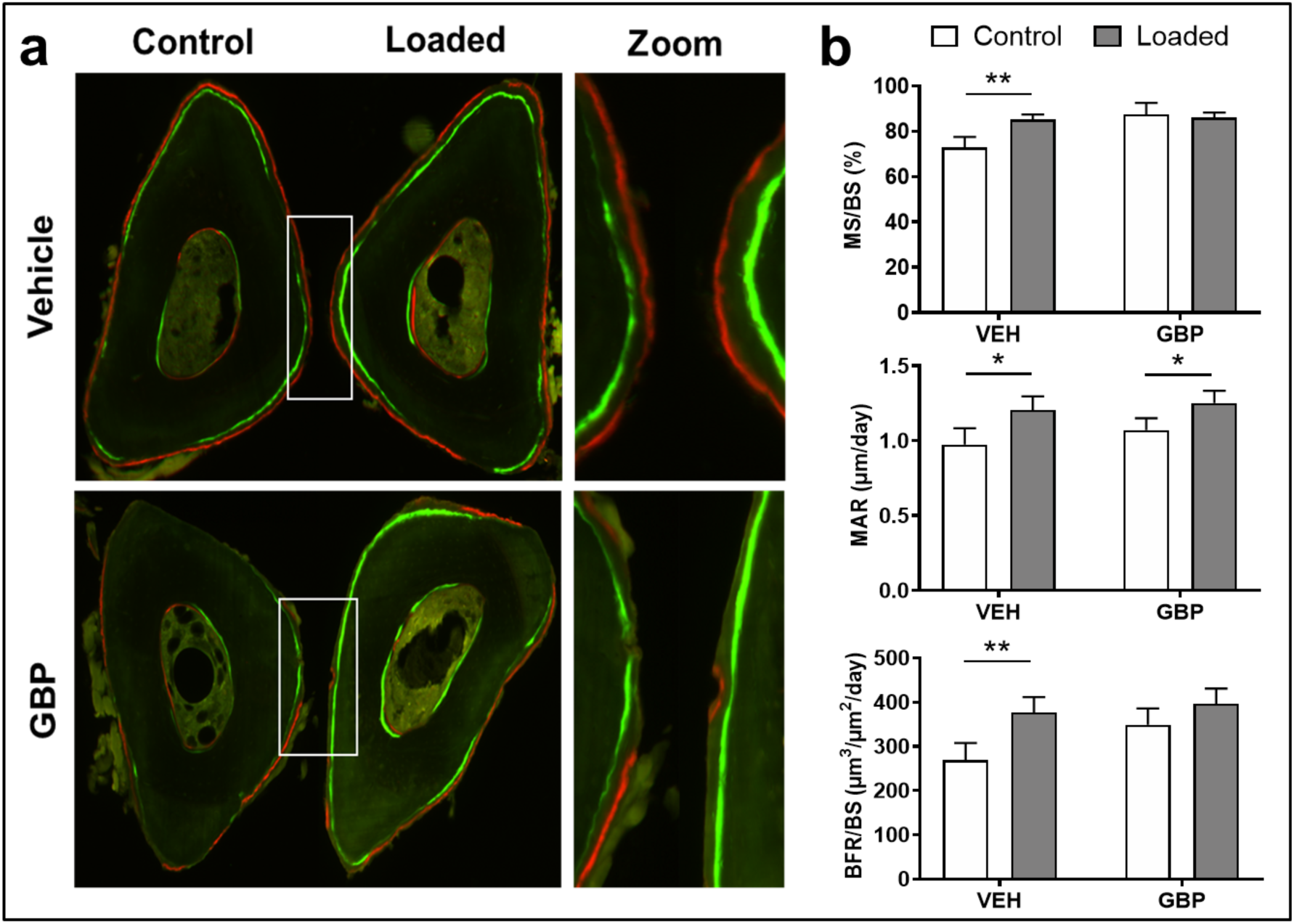
Gabapentin treatment decreases ulnar mechanosensitivity and load-induced bone formation. Male C57BL/6J mice were injected daily with saline (vehicle, VEH) (n=9) or Gabapentin (GBP, 300mg/kg BW) (n=7) for 4 weeks while undergoing axial ulnar loading. **(a)** Representative images of control (non-loaded) and loaded ulnas from VEH and GBP treated mice. Changes in **(b)** mineralizing surface (MS/BS), mineral apposition rate (MAR), and bone formation rate (BFR/BS) in response to mechanical loading were assessed in VEH and GBP treated mice. Paired Student’s t tests compared control, contralateral ulnas to loaded ulnas. Values are shown as Mean ± SEM; p≤0.01(**), p≤0.05(*).

**Table 3.**
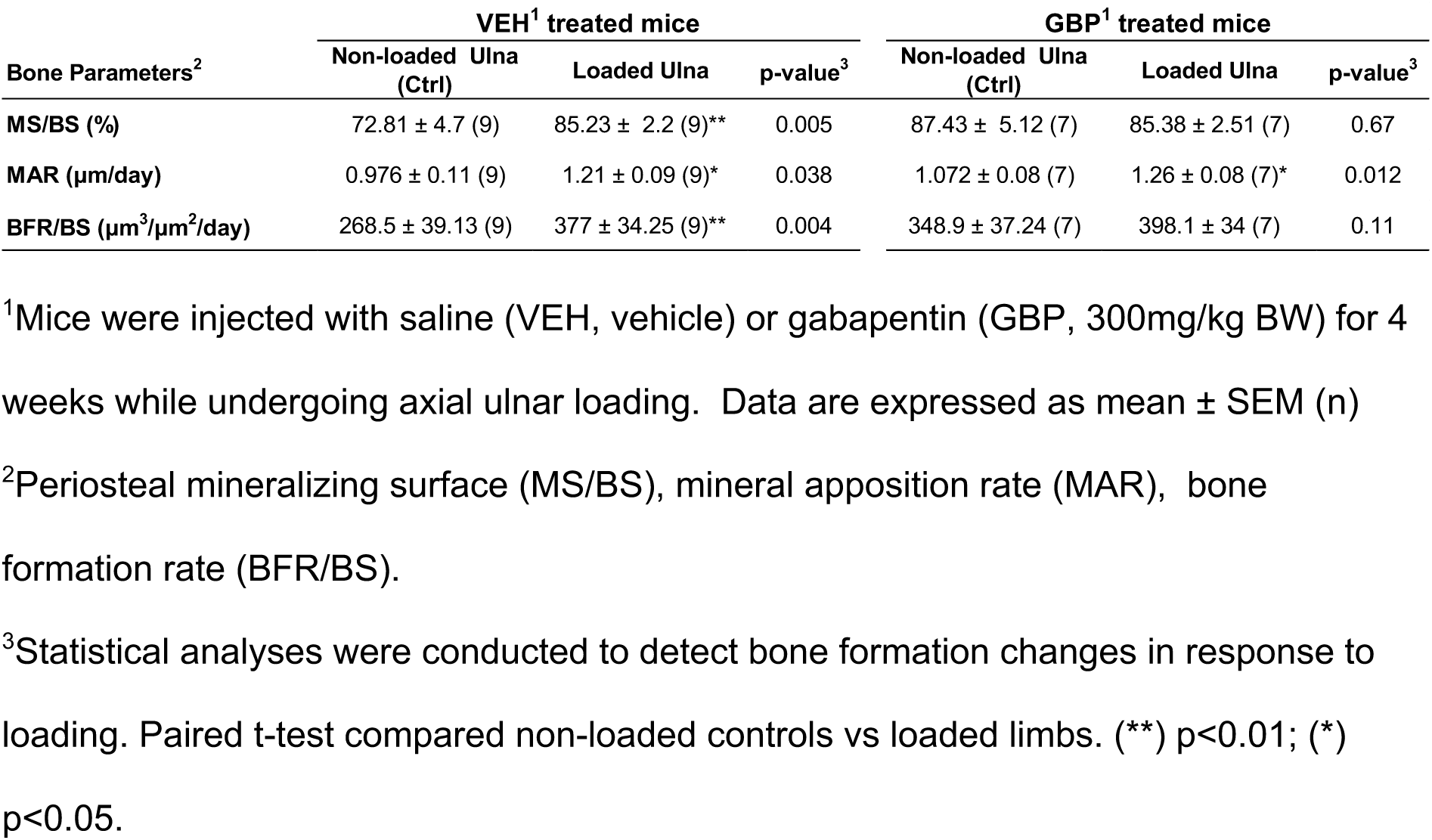
Ulnar dynamic histomorphometry measurements in response to mechanical loading in vehicle and gabapentin treated mice

## Discussion

Mechanotransduction requires physical coupling of mechanosensory components and the ability of those components to transduce mechanical signals into biochemical responses^29^. Numerous studies have identified molecules that contribute to mechanical signaling within bone such as sclerostin^30, 31^, connexins^32–34^, and focal adhesions^35–39^. However, the mechanism by which force is directly transmitted from the bone matrix to the osteocyte cell membrane remains unclear. Likewise, while the presence of transverse TEs in osteocytes has been established^11, 13, 24, 40^, the cell membrane molecules to which PLN-containing tethers bind is unknown.

Our hypothesis that PLN binds to the α_2_δ_1_ subunit of VSCCs was formed through several observations. First, various studies showed that VSCCs regulate skeletal mechanosensitivity^16, 41, 42^. Second, spatial positioning of α_2_δ_1_ is optimal for interaction with PLN, in that α_2_δ_1_ has a large extracellular region (α_2_) capable of interacting with ligands. And third, the ability of α_2_δ_1_ to regulate gating kinetics of the α_1_ pore of VSCCs^43^ made α_2_δ_1_ a strong candidate receptor for PLN binding. Our data showed that PLN matrix tethers bind α_2_δ_1_ with high affinity, connecting the mineralized bone matrix with the osteocyte cell membrane (**Fig. 5a**).

**Figure 5.**
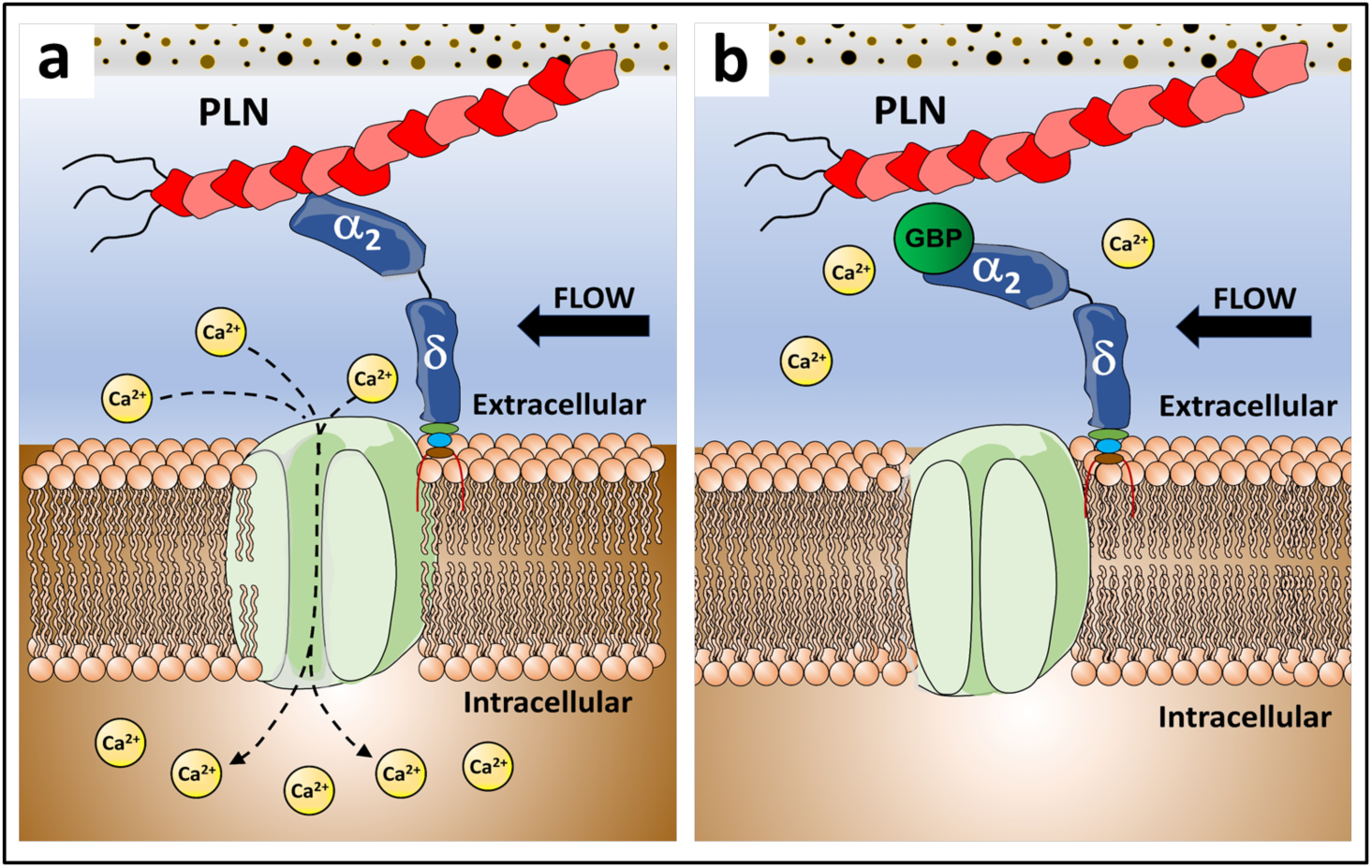
Summary of results. In this work we found that **(a)** the α_2_δ_1_ subunit of voltage sensitive Ca^2+^ channels binds perlecan (PLN) creating a mechanosensory complex that enables connection between the mineralized matrix and the osteocyte cell membrane. **(b)** We also demonstrated that gabapentin (GBP) interferes with binding of PLN and α_2_δ_1_ *in vitro*. As the PLN/α_2_δ_1_ complex is necessary for mechanotransduction, GBP uncoupling of the complex results in impaired osteocyte mechanosensation *in vivo*, which may account for the deleterious skeletal effects observed with chronic use of this drug.

We previously demonstrated that α_2_δ_1_ modulates mechanically-regulated ATP release in osteocytes via its association with Ca_v_3.2 (α_1H_), the predominant α_1_ pore-forming subunit within these cells^24, 44^. The extracellular portion (α_2_) of the α_2_δ_1_ subunit is known to be glycosylated with N-acetyl glucosamine sugars. These glycosylation sites are essential for surface expression of α_2_δ_1_ and have high affinity to WGA^45^. In this work we confirmed expression of α_1H_ in MLO-Y4 cells and found that PLN staining independently overlapped with α_1H_ and WGA fluorescent signals at the cell surface of osteocytic cells, suggesting close physical proximity of PLN and the α_1H_ pore and the sugars attached to α_2_δ_1_. In addition, α_2_δ_1_ and PLN co-localize in osteocytic cells along the dendritic processes of osteocytes, the area most sensitive to mechanical force^46^. Furthermore, by quantifying the molecular interactions between the extracellular portion of α_2_δ_1_ and different PLN domains/subdomains, we demonstrated that α_2_δ_1_ and PLN binding is facilitated within the cysteine-free region of PLN Dm III-2, with *K_D_* values in the low nanomolar range compared to other PLN subdomains, showing *K_D_* values in the milli- and micromolar ranges. As a reference, binding of biotin and avidin is among the strongest non-covalent affinities known^47^ with a dissociation constant of about 1.3 x 10^-15^. This aligns with literature reports in which Dm III mediates the binding of other molecules with PLN, including the fibroblast growth factor (FGF)-7 (N-terminal half of Dm III)^48^, platelet-derived growth factor (PDGF) (Dm III-2)^49^, and FGF18 (Dm III, cysteine-free region)^50^. Further, previous work showed that PLN binds to another matrix molecule called von Willebrand Factor A-domain-Related Protein (WARP)^51^. Notably, the interaction between WARP and PLN is mediated through Dm III-2 of PLN and the von Willebrand Factor A (vWFA) domain of WARP^51^. As the α_2_ portion of the α_2_δ_1_ subunit contains a vWFA domain^52^ which enables binding to extracellular matrix molecules^53^, these findings provided further reasoning that PLN and α_2_δ_1_ form a functional complex.

*In silico* docking models between the vWFA domain of α_2_ and PLN Dm III-2 predicted strong interactions between these molecules. Although there are limitations in the interpretation of HDOCK results^27, 28^, the quality results for structure accuracy indicate that the docking predictions obtained are reliable. Our 3D models, combined with the LSPR data, confirm that Dm III-2 is a binding site for α_2_, mediating the interaction of the PLN/α_2_δ_1_ complex. Together, this M-CTC, composed of PLN and α_2_δ_1_, is thus spatially, structurally, and biochemically positioned to activate osteocytes in response to mechanical force (**Fig. 5a**).

Whereas several clinical studies link chronic use of GBP with adverse skeletal side effects, including increased fracture risk^22, 54^, the molecular mechanisms underlying these effects and whether they occur directly in bone are entirely unknown. We hypothesized that GBP disrupts PLN/α_2_δ_1_ binding, affecting the function of the M-CTC, which may explain the skeletal side effects of this medication. GBP recognizes an Arg-Arg-Arg (RRR) motif within the α_2_ region of the α_2_δ_1_ subunit^20^, located upstream and in close proximity to the vWFA domain **(Fig. 1)**. Interactions occurring in regions flanking the vWFA can restrict the conformation of the domain (i.e., close, low affinity vs open, high affinity ligand binding states)^55^. Thus, binding of GBP to the RRR motif may disrupt vWFA-mediated interactions of α_2_δ_1_ with other proteins, such as was demonstrated in a recent study where GBP blocked binding of α_2_δ_1_ and thrombospondins^56^. Here, we demonstrated that GBP interferes with binding of PLN (full-length and Dm III-2) and α_2_δ_1_ *in vitro*, effectively uncoupling the M-CTC (**Fig. 5b**). We also showed that acute GBP treatment in mice blunts the anabolic bone responses to mechanical loading. Previous studies have shown that both PLN^13^ and α_2_δ_1_ (ref. 24) are necessary for mechanotransduction in skeletal cells. Thus, GBP may impair osteocyte mechanosensation by disrupting the function of the PLN::α_2_δ_1_ complex and contribute to the deleterious skeletal effects observed with chronic use of these drugs^21–23^.

Limitations of this study include the use of only male mice for evaluating the *in vivo* effects of GBP on bone. Ongoing work is focused on understanding the tissue level impact of GBP in female mice. Furthermore, we did not assess binding of PLN Dm II with α_2_δ_1._ However, as we found that Dm III-2 bound with equivalent affinity to that of full-length PLN, we were confident that the observed binding between the full-size core protein of PLN was mediated through Dm III-2. Additionally, in contrast to that of PLN Dm III and α_2_δ_1_, there are no previous studies that support a potential interaction between Dm II and α_2_δ_1._

Notable strengths of this work included the use of LSPR-based experiments to determine the interactions between PLN and α_2_δ_1_. In this regard, the nanoplasmonic sensors provided reproducible limit of detection at the low zeptomolar range, along with quantitative dissociation constant values (*K_D_*) between biomolecules^57, 58^ with far greater sensitivity than conventional SPR methods. Additionally, while our interest in the M-CTC lie in osteocyte physiology, it is likely that the function of this complex is conserved across numerous tissues. As such, identification of this novel mechanosensory complex may have a dramatic impact on understanding how other tissues regulate mechanosensation, especially as PLN serves mechanotransduction functions in other cell types^59^.

In summary, this work identified novel interactions between the large heparan sulfate proteoglycan PLN and an extracellular auxiliary subunit of VSCCs. Formation of this complex revealed how the transverse tethers previously identified as force transducers in osteocytes attach to the cell membrane, but also provided a greatly expanded understanding of how VSCCs are capable of being activated by mechanical force. Most importantly, our data demonstrate how GBP may negatively regulate bone remodeling by interfering with osteocyte mechanosensation. Better understanding of the mechanisms by which GBP regulates skeletal mechanotransduction will guide the treatment of patients using these drugs and may lead to the design of precision agents efficacious at their target tissues, but devoid of detrimental skeletal effects.

## Materials and Methods

### Cell culture and immunofluorescence

Immunofluorescence experiments were performed using the osteocytic cell line MLO-Y4. Approximately 1,000 MLO-Y4 cells were seeded onto collagen-I coated 8-well chambers (NUNC™, Rochester, NY) and cultured as described previously^60^. When cells were 80-90% confluent, media was removed, cells were washed with Tris-buffered saline (TBS) and fixed with paraformaldehyde (4%, v/v) diluted in TBS for 45 min at room temperature (RT). Cells were washed with TBS to remove residual fixative and incubated for 1h at RT in donkey serum (5%, v/v) diluted in TBS with Tween 20 (0.1 %, v/v). Cells were incubated with the appropriate primary antibodies (Abs) diluted in blocking buffer for 1h at RT. For co-localization experiments, where association between Ca_v_3.2 (α_1H_), α_2_δ_1,_ and PLN were performed in osteocytic cells, the following primary Abs were used: affinity-purified rabbit anti-Ca_v_3.2 (α₁_H_) polyclonal antibody (1:100) was raised against a synthetic peptide sequence and prepared for our laboratory commercially by ResGen (Invitrogen, Carlsbad, CA, USA), as described^61^. Staining for α_2_δ_1_ was performed as previously reported^24^, affinity purified rabbit anti-α_2_δ_1A_ isoform polyclonal antibody (1:500) produced by Bethyl Laboratories (Montgomery, TX)^62^ was used. For PLN staining, cells were incubated with rat monoclonal anti-PLN domain-IV (A7L6) primary antibody (1:40) (Abcam, Boston, MA, USA). Following incubation with the primary Abs, cells were washed with blocking solution and incubated with species-specific Alexa Fluor 488 and 555 conjugated secondary Abs (1:200) (Invitrogen, Carlsbad, CA, USA) and DR™ nuclear stain (1:1000) (Biostatus, Ltd, Shepshed Leicestershire, UK) diluted in blocking solution. To visualize cell membrane glycoproteins, cells were stained with fluorescein conjugated wheat germ agglutinin (WGA) (Invitrogen, Carlsbad, CA, USA). Samples were washed with TBS, mounted, and stored at 4°C until imaged. Negative controls for cultured cells were performed using non-immune IgGs diluted at concentrations equivalent to primary Abs or without primary Abs. For cells stained with WGA, an N, N’, N’’-triacetylchitotriose control was used. Samples were imaged with an LSM 510 VIS confocal microscope using a 40X C-apochromat water immersion objective (NA 1.2) (Zeiss, Inc, Thornwood, NY).

### Co-immunoprecipitation and western blotting

To determine if α_2_δ_1_ associates with PLN, co-immunoprecipitation assays were performed. MLO-Y4 cells (∼90% confluent) cultured on 100 mm dishes were exposed to 500 μL of radio immunoprecipitation (RIPA) lysis buffer containing a protease inhibitor cocktail added just prior to cell lysis (1:100) (Sigma-Aldrich, USA). Plates were incubated with lysis buffer at 4°C for 1 min. Lysates were scraped from each plate and placed in 1.5 mL tubes. Samples were sonicated and centrifuged (14,000 g) for 10 min at 4°C. Protein concentration was determined using the Pierce BCA protein assay kit (ThermoFisher Scientific, MA, USA). Samples were diluted in RIPA buffer to achieve equal protein concentrations. Pre-cleared lysates were added to 100 μL of magnetic Dynabeads (Invitrogen, Carlsbad, CA, USA) complexed to 5 ug of monoclonal anti-PLN A7L6 antibody (Abcam, Boston, MA, USA) or Rat IgG. Lysates and beads were incubated on a rotator at RT for 45 min. The Dynabead-Ab-Ag complex was washed three times with 1X phosphate-buffered saline (PBS). Beads then were resuspended in PBS and the supernatant was transferred to a new tube. Supernatants were diluted in Laemmli buffer containing β-mercaptoethanol (2%, v/v) and boiled for 10 min. Western blotting was performed as described^38^. Equal volumes of each sample (20 µL) were electrophoresed in 8-12% Tris-Acetate gels and probed with the anti-α_2_δ_1A_ (Bethyl Laboratories) and anti-PLN A7L6 (Abcam, Boston, MA, USA) primary antibodies (1:500). Blots were probed for β-actin Ab (Cell signaling) (1:500) as a loading control.

### Recombinant α_2_δ_1_ polypeptides

The α_2_ portion of the human α_2_δ_1_ protein (NCBI reference sequence NP_00713.2) was produced by *GenScript Protein Expression and Purification Services* (GenScript Corp, Piscataway, NJ). Briefly, the α_2_ target DNA sequence was designed, optimized, and synthesized by sub-cloning into a pcDNA3.4 vector and transfection-grade plasmid was maxi-prepared for cell expression. Expi293F cells were grown in serum-free Expi293FTM Expression Medium (ThermoFisher Scientific, MA, USA). Cells were maintained in Erlenmeyer flasks (Corning, NY, USA) at 37°C with CO_2_ (8% v/v) on an orbital shaker (VWR Scientific). One day before transfection, cells were seeded at an appropriate density in flasks. On the day of transfection, DNA and transfection reagent were mixed at an optimal ratio and added to the cells. The recombinant plasmid encoding the target protein was transiently transfected into Expi293F cells. Culture supernatants, collected on day 6, were used for protein purification. Conditioned media was centrifuged, filtered, then passed through a HisTrapTM FF crude affinity purification column at an appropriate flowrate. After washing and elution with appropriate buffers, the eluted fractions were pooled, and buffer exchanged to the final formulation buffer. Purified protein was analyzed by western blot to confirm the molecular weight and purity. The concentration was determined by Micro-Bradford assay with BSA as a standard (ThermoFisher Scientific, MA, USA). Purified protein was stored in 1x PBS (pH 7.2), filter sterilized (0.22 μm), and packaged aseptically at a concentration of 37 μg/mL.

### Full-length perlecan and perlecan domains I, III, IV-1, IV-2, IV-3, and V

Full-length PLN was isolated and purified from HT-29 human colorectal cancer cells (formerly called WiDr) (ATCC, Manassas, VA, USA) as reported^63, 64^ (**Suppl. Methods**). PLN domains (Dm) I, Dm IV-1, Dm IV-2, and Dm IV-3 (ref. 63, 65, 66) and Dm V (ref. 67), were produced and purified as described previously (**Suppl. Methods**). PLN Dm-III is composed of three cysteine-free, laminin-like globular domains with alternating laminin EGF-like cysteine-rich regions^68^. We designed two Dm-III plasmids using SnapGene, the first encoding the cysteine free, globular region of PLN Dm III-2 (laminin IV-A2) and the second Dm III-2 (IV-A2) followed by a cysteine-rich laminin EGF-like region (Dm III-2 + cysteine). Each contained an EF-1α promoter and BM40 signal sequence for enhanced secretion, as well as a C-terminal FLAG tag and 6x His-tag for purification (VectorBuilder, IL, USA). Plasmids were transfected into HEK293A cells using Lipofectamine 2000 (Life Technologies, CA, USA). Transfected cells were grown from single-cell clones and selected with G418 (2 mg/mL). Dm III-2 and Dm III-2 + cys production was confirmed via western blot using 6x His-tag Ab (Invitrogen, Carlsbad, CA, USA). Positive clones were expanded, purified, and sequenced for verification. Conditioned media from hyperflasks was collected and concentrated in bulk using the Sartorius Vivaflow Cross-flow System (Sartorius, NY, USA) with Vivaflow 200 10,000 MWCO PES filters (Sartorius, NY, USA). Dm III-2 and Dm III-2 + cys were purified using Ni-NTA resin as described for Dm IV recombinant proteins **(Suppl. Methods)** with one additional wash of 500 mM NaCl after conditioned medium flow through and before the imidazole (20 mM) wash. The purified protein was buffer exchanged and stored at - 80°C.

### Localized surface plasmon resonance (LSPR) experiments

The LSPR-based assay was used to delineate the region of each protein necessary for the structural integrity of the matrix-channel tethering complex (M-CTC), which enabled quantification of the binding interaction between full-length PLN, recombinant subdomains of PLN, and the α_2_ portion of the α_2_δ_1_ subunit. In brief, noble metal nanoparticles display unique localized SPR properties, which are dependent on the size and shape^69–71^, and most importantly, the dielectric constant of their surrounding environment^72, 73^. Utilizing the latter dependency, solid-state, LSPR-based sensors have been developed employing simple optical spectroscopy to detect biological constituents by monitoring the LSPR changes (ι1α) induced by their presence^58, 74^. A schematic representation of LSPR experiments is summarized in **Supplementary Figure S2. *Synthesis of gold triangular nanoprisms (Au TNPs)*.** Au TNPs were chemically synthesized according to published procedures^75–77^. Briefly, 10.4 mg (0.05 mM) of Et_3_Pau(I)Cl were dissolved in N_2_ purged acetonitrile (20 mL) and stirred at RT for 5-10 min. Then, 19 µL (0.273 mM) of triethanolamine (TEA) was added to the solution and heated. Upon solution temperature reaching 38 °C, 300 µL of polymethylhydrosiloxane (PMHS) was added, and the reaction slowly stirred. During the reaction, the solution color changed gradually from colorless to dark navy-blue, indicating the formation of Au TNPs. Once a dark navy-blue color was achieved, the LSPR dipole peak position (λ_LSPR_) was monitored through UV-visible spectroscopy until the solution was λ_LSPR_ = ∼800 nm, indicating the formation of ∼42 nm edge length Au TNPs (***SI appendix,* Fig. S3**). The Au TNP solution was centrifuged at 7,000 rpm for 10 s, transferred to 3-mercaptopropyltrimethoxysilane (MPTMS)-functionalized glass coverslips (***SI appendix,* Suppl. methods**) and incubated for 1 h. TNP bound coverslips were rinsed with acetonitrile, dried with N_2_ gas, and stored under N_2_ at 4°C. Au TNP-bound coverslips were used within 3 days of the attachment**. *α_2_-functionalized Au TNPs*.** Au TNP-bound glass coverslips underwent a tape-cleaning procedure to remove non-prismatic structures. Briefly, 3M adhesive tape was placed onto the Au TNP-bound glass coverslip, pressed gently with the thumb, and then the tape was removed at a 90^0^ angle. Cleaned Au TNP-bound coverslips were cut into 6.25 mm x 25 mm pieces using a diamond cutter to produce the sensors (**Suppl. Fig. S2a**). Each sensor was incubated in 6.0 mL of a 1.0 mM:1.0 µM ratio of 11-mercaptoundecanoic acid (MUDA): 1-nonanethiol (NT) solution overnight (**Suppl. Fig. S2b**). The following day, the sensors were rinsed with ethanol to remove loosely bound thiols. This thiol treatment created a self- assembled monolayer (SAM) onto Au TNP surface. Next, SAM-modified Au TNPS were incubated in an EDC/NHS (0.2 M) solution for 2 h to activate the acid group of MUDA, rinsed with ethanol and PBS, and incubated overnight in a PBS buffer solution (pH 7.2) containing the α_2_ portion of α_2_δ_1_ (10 ng/mL) (**Suppl. Fig. S2c**). To determine the dissociation constant (*K_D_*) values for interactions between α_2_ and PLN, each α_2_-functionalized sensor was rinsed with PBS and incubated in a solution containing different concentrations (1x10^-16^ to 1x10^-8^ M) of full-length PLN (digested with heparanase and chondroitinase, or undigested) or each of PLN domains/subdomains Dm I, III-2 (cys free), III-2 (cys), IV-1, -2 and -3 or V (**Suppl. Fig. S2d**). At the end of the experiments, the sensors were removed for data collection. Once we established the regions of PLN that mediate binding within the M-CTC, assays were repeated with the addition of GBP (see *drug binding experiments*). ***Protein binding curves and spectroscopy characterization*.** Before and after each incubation step, an extinction spectrum of the sensor was collected through UV-visible spectroscopy, and the shift in the LSPR dipole peak position (*Δλ*_LSPR_) was obtained (**Suppl. Fig. S2e**, **S3**). All absorption and extinction spectra were collected utilizing a Varian Cary 50 Scan UV-visible spectrometer in the range of 300-1,100 nm, using 1 cm quartz cuvettes. All spectra were collected in ethanol or PBS (pH 7.2) to keep the bulk refractive index constant. The “background” was a coverslip immersed in ethanol/PBS. The reference (blank) was a sensor incubated in ethanol/PBS (no analyte present). Scanning electron microscopy (SEM) images of Au TNPs were characterized using a JEOL 7800F SEM. ***Data Processing*.** For all UV-vis extinction spectra, *λ_L_*_SPR_ was determined through curve fitting using OriginLab software. The *Δλ*_LSPR_ was calculated by taking the difference between the λ_LSPR_ before and after each fabrication step. *Δλ*_LSPR_ values were reported as the Mean ± standard deviation (SD) of six individual measurements at each concentration used. Using the statistics software GraphPad Prism, protein binding curves were developed by plotting *Δλ*_LSPR_ versus PLN [or PLN subdomains] concentration in mol/L (M) (**Suppl. Fig. S2f**). Binding curves were fitted to a specific binding Hill slope (**Suppl. methods**) to determine the *K_D_* values between α_2_ and PLN domains/subdomains.

### Drug binding experiments

LSPR-based experiments were also used to determine the interactions among α_2,_ PLN, and GBP. Three different approaches were performed. First, to determine if GBP was capable of displacing PLN from α_2_ following binding of PLN to α_2_, the α_2_-functionalized sensors were incubated overnight with full-length PLN (10 nM) or PLN Dm III-2 (100 nM), followed by overnight incubation with GBP (0.33 mg/mL).

Second, to determine if PLN could displace GBP from α_2_, the α_2_-functionalized sensors were incubated overnight with GBP (0.33 mg/mL), followed by overnight incubation with full-length PLN (10 nM) or PLN Dm III-2 (100 nM). Lastly, to determine if PLN or GBP had greater affinity for α_2_ when provided equal opportunity to bind, the α_2_-functionalized sensors were incubated overnight in a mixture of full-length PLN (10 nM) or PLN Dm III-2 (100 nM) and GBP (0.33 mg/mL). At the end of the experiments, sensors were removed for data collection and processing as described above.

### *3D* docking models

*In silico* protein-protein, functional interactions and 3D docking models between the vWFA domain of α_2_δ_1_ and domain III-2 of PLN were simulated with the free web HDOCK^27, 28^ server (http://hdock.phys.hust.edu.cn/). To develop high confidence homology models of protein structures, multiple sequence alignment was conducted using Clustal Omega (1.2.4) (ref. 78) (https://www.ebi.ac.uk/Tools/msa/clustalo/). For PLN, the sequences of the three Laminin-IV A subdomains in PLN Dm III [P98160 residues 538-730 (Dm III-1); 941-1125 (Dm III-2), and 1344-1529 (Dm III-3)] were aligned first. Then, the sequence of PLN Dm III-2 [P98160, residues 941-1125] was selected to be aligned against the sequences of the Laminin IV type A1 (P24043; residues 531-723) and Laminin IV type A2 (P24043; residues 1176-1379) domains of Laminin alpha-2. For the vWFA domain, the sequences of the vWFA domains of human thrombospondin 1 (P07996; residues 316-373), thrombospondin 2 (P35442, residues 318-375) and α_2_δ_1_ (residues 253–430 of CACNA2D1 [P54289]) were used for ClustalO alignment. The amino acid sequences for the vWFA domain of the α_2_ peptide (residues 253–430 of CACNA2D1 [P54289]) and PLN Dm III-2 (residues 941-1125 of HSPG2 [P98160]) were input into the protein fold recognition server Phyre2 (ref. 79) to obtain structural 3D models using known protein templates. The structures with the higher model confidence (the probability that the match between the input sequence and the template is a true homology) and I.D. value (the percentage identity between the input sequence and the template) were chosen for docking. The protein template information and 3D structures were retrieved from the RCSB protein data bank (https://www.rcsb.org/). For the vWFA domain of α_2_δ_1_, the structure of the von Willebrand factor type A from Catenulispora acidiphila (4FX5) (https://www.rcsb.org/structure/4FX5) was selected. For PLN Dm III-2, the structure of the L4b domain of human Laminin alpha-2 (4YEP) (ref. 80) (https://www.rcsb.org/structure/4YEP) was used as the best match. In the HDOCK server, PDB files for 4FX5 (vWFA) and 4YEP (PLN Dm III-2) were used to populate the information for receptor and ligand, respectively. The output with the highest docking energy score from the top 10 predictions was selected for visualization.

### Animal experiments and in vivo ulnar loading

Male C57BL/6J mice were purchased from the Jackson Laboratory (JAX, Bar Harbor, Maine) and group-housed (2–4 mice/cage) on TEK-fresh bedding in ventilated cage systems at the Indiana University School of Medicine animal facilities. Food and water were provided *ad libitum* and mice were maintained under 12-h light/dark cycles and standard conditions of temperature and humidity. At 16 weeks of age, mice were randomly assigned into 2 groups to receive daily intraperitoneal injections of saline (vehicle, VEH) or gabapentin (GBP, 300mg/kg BW; 50mg/mL stock diluted in saline) (Acros Organics AC458020050, ThermoFisher Scientific, MA, USA) for 4 weeks (n=9 mice/treatment). Sample size calculations were based on published data to detect histomorphometrically-measured changes in bone formation induced by loading of 100 µm^3^/µm^2^/yr, and a true difference between loaded and non-loaded bones as small as 40 µm^3^/µm^2^/yr (α=0.05 level; power (1–β) = 80%).

GBP and VEH treated mice were subjected to axial ulnar compression to induce anabolic skeletal responses as previously described^30^. Briefly, mice were anesthetized under gas isoflurane and the right ulna was loaded using a sinusoidal (haversine) waveform (-2200 µε, 2 Hz, 180 cycles). Mice received one loading bout every other day over a 10-day period, loading order of mice was randomized each time. Left ulnas were used as non-loaded, contralateral controls. To monitor load-induced bone formation, the fluorochromes calcein (10 mg/kg, Sigma-Aldrich, USA) and alizarin (20 mg/kg, Sigma-Aldrich, USA) were injected intraperitoneally one day before the final loading bout and 11 days later, respectively. Mice were euthanized at 20 weeks of age by CO_2_ asphyxiation, followed by cervical dislocation. Ulnas were harvested and processed for dynamic histomorphometry as published^30^. All experiments conducted were approved by the Indiana University Institutional Animal Care and Use Committee.

### Dynamic histomorphometry

Preparation and histological sectioning of ulnas was conducted by the Histology and Histomorphometry Core within the Indiana Center for Musculoskeletal Health at Indiana University. To detect bone formation changes in double-labeled histological sections, the following parameters were assessed as previously described^30^: periosteal mineralizing surface (MS/BS, %), mineral apposition rate (MAR, μm/day), and bone formation rate (BFR/BS, μm^3^/μm^2^/day). All measurements were collected such that investigators were blinded to treatment.

Statistical analyses were conducted using the GraphPad Prism software version 9.3.1(471) (La Jolla, CA). Paired Student’s t tests compared control, contralateral ulnas to loaded ulnas. Results are reported as mean ± standard error of the mean (SEM). Significance level was defined as p≤0.05.

### Data availability

All data generated or analyzed during this study are included in this published article and its supplementary information files.

## Acknowledgements

This study was supported by 1R01AR074473-01 and UL1 TR001108 to WRT, Indiana University Research Support Funds Grant to RS and WRT, 1F32AR074893-01 to CSW, and Faculty Research Development funds through Marian University to JMH and WRT.

## Author Contributions

PCRF, XY, ANM, TVT, AB, WRT collection/assembly of data. PCRF, XY, ANM, TVT, AB, CSW, KR, GB, DW, AGR, KR, MLN, KEW, KJL, US, JMH, RS MCFC, WRT data analysis/interpretation. MLN, KEW, KJL, US, JMH, RS, MCFC, WRT concept/design. PCRF, MCFC, WRT manuscript writing. All authors have seen and approved the submitted manuscript.

## Supplementary Information

Supplementary information accompanies the manuscript on the Bone Research website http://www.nature.com/boneres

