## Supplemental information for "Gabapentin Disrupts Binding of Perlecan to the α_2_δ_1_ Voltage Sensitive Calcium Channel Subunit and Impairs Skeletal Mechanosensation"

**This PDF file includes:**

Supplementary Methods
Supplementary Figures S1 to S4
Supplementary Table S1

### Supplementary Methods

#### Perlecan recombinant proteins

***Full-length perlecan***. Full-length PLN was isolated and purified from HT-29 human colorectal cancer cells (formerly called WiDr) (ATCC, Manassas, VA, USA) as reported[1,2]. Briefly, cells were cultured and maintained in Eagle's minimal essential medium (EMEM) (ATCC® 30-2003), fetal bovine serum (FBS) (10%, v/v) and penicillin/streptomycin (1% v/v). When cells were 100% confluent, fresh media with lower FBS levels (2%, v/v) was added and conditioned medium was collected. The conditioned medium was filtered and concentrated using the Amicon stirred cell pressure-driven filtration device (400 mL, 46 mm membrane filter) (Millipore, Sigma). The resulting high molecular weight concentrated solution was subjected to anion exchange chromatography containing positively charged diethylaminoethyl (DEAE)-sepharose, facilitating binding of negatively charged heparan sulfate chains. The DEAE column employed measured 2.5 cm x 10 cm. A 50 mL bed volume was equilibrated with a buffer containing: Tris/HCl (0.05 M, pH 8.6), urea (2 M), NaCl (0.25 M), EDTA (2.5 mM), benzamidine (0.5 mM), PMSF (0.5 mM) and sodium azide (0.02%, w/v). Four volumes of equilibration buffer were passed through the column before the concentrated media was loaded. Concentrated media was passed over the equilibrated column at 4ºC 10 times until all 500 mL of media were loaded. The column then was washed with 4 volumes of equilibration buffer at a flow rate of approximately 1 mL/min, collecting 1.5-2.0 mL fractions. Fractions were followed using absorbance readings at 280 nm. Bound molecules then were eluted using a high salt buffer (same as equilibration buffer but with 0.75 M NaCl). Eluants then were separated by size-exclusion chromatography using a column containing Sepharose CL-4B. Fractions were analyzed by dot blot immunoassay using anti-PLN (A7L6) Ab. Pooled fractions containing PLN were dialyzed in dd$H_2O$ and centrifuged in a speed vac until the desired concentration was obtained. Enriched PLN solution was separated using HiPrep Heparin FF affinity column to bind heparan sulfate chains and subsequently obtaining purified PLN protein. Enriched fractions were pooled, concentrated, and dialyzed into a final PBS working solution. Purity was assessed by silver stain and samples were aliquoted and stored at −80°C until use.

***Perlecan Domains I, IV-1, IV-2 and IV-3.*** PLN domain (Dm) I, Dm IV-1, Dm IV-2, and Dm IV-3, were produced as described previously[1,3,4]. Briefly, transfected HEK293A cells expressing recombinant proteins were expanded and cultured in hyperflasks (Corning, NY, USA) in Dulbecco's Modified Eagle Medium (DMEM) (Corning, NY, USA) with FBS (2%, v/v), penicillin/streptomycin (1%, v/v), and either puromycin (400 ng/mL) (ThermoFisher Scientific, MA, USA) for Dm IV-1 and IV-2, or Geneticin (G418) (1 mg/mL) (Teknova, CA, USA) for Dm I and IV-3. Full media exchanges were conducted every three days. Inhibitors phenylmethylsulfonylfluoride (PSMF, 0.5 mM) and benzamidine (0.5 mM), and sodium azide (0.02%, w/v) were added to conditioned media and filtered through a PES filter flask (0.2 µm, Corning, NY, USA). Conditioned media was concentrated using a 10,000 Dalton cut-off filter then passed through Ni-NTA agarose resin (ThermoFisher Scientific, MA, USA). The column was first equilibrated with imidazole (10 mM) in PBS, conditioned media was applied, then the column was washed with imidazole (20 mM) in PBS, then eluted with imidazole (300 mM) in PBS. Eluted fractions were pooled, and buffer exchanged with Millipore Amicon Ultra-15 Centrifugal Filter Units (Millipore Sigma, MA, USA).

***Perlecan Domain V***. PLN Dm V was produced and purified as previously described[5]. Human Dm V was cloned into the pSecTag2A vector (Invitrogen, Carlsbad, CA, USA) using the primers 5′ Dm V Asci pSecTag2A: 5′-AGGGCGCGCCATCAAGATCACCTTCCGGC-3′; 3′ Dm V Xho1 pSEcTag2A: 5′-AGCTCGAGCCGAGGGGCAGGGGCGTGTGTTG-3′. To confirm that Dm V effects were not due to any single clone-specific irregularities, Dm V was also cloned into the vector pCepPu (provided by Maurizio Mongiat, Center for Cancer Research, Aviano, Italy) using the following primers: NHEI whole Dm V forward 5′-AGGCTAGCGATCAAGATCACCTTCCGGC-3′; XHOI HIS Dm V reverse 5′-AGCTCGAGCATGATGATGATGATGATGCGAGG-3′. The Dm V cDNA was amplified from human umbilical vein endothelial cell (HUVEC) cDNA using a GC-rich PCR system and dNTPack (Roche Applied Science). Maxi-preps of Dm V DNA were transfected into 293FT (for pSecTag2A vector, ATCC) or 293 EBNA (for pCepPu vector) cells via Lipofectamine (Invitrogen, Carlsbad, CA, USA). After transfection, the 293 cells were put into a CELLine Adhere 1000 bioreactor (Argos Technologies) and grown for 7 days in complete medium containing FBS (10%, v/v), antibiotic/antimycotic, G418 sulfate (1%, v/v), and puromycin (0.05 µg). After 7 days the complete medium was removed. Cells then were washed 5 times with CD293 medium containing L-glutamine (4 mM), penicillin/streptomycin (1%, v/v), G418 sulfate (1%, v/v), and puromycin (0.05 µg) to remove any serum; and then fresh CD293 medium was added to the cells. Cells then were incubated for 7 days, followed by collection of Dm V-containing conditioned medium, and purification of Dm V via its C-terminal 6X His-tag and Ni-ATA agarose beads (QIAGEN) per manufacturer's protocol. Eluted fractions containing Dm V were combined and dialyzed against PBS and the purity of the resultant Dm V was confirmed via SDS-PAGE stained with Brilliant Blue G Colloidal, silver stain (FASTsilver Gel Staining Kit, Calbiochem) following the manufacturer's protocol.

**Localized surface plasmon resonance experiments**

***Silanization of glass coverslips.*** Glass coverslips (25 x 25 mm) were silanized as previously described[6-9]. Briefly, glass coverslips were placed in a glass staining jar, incubated in a 10% RBS 35 detergent solution at 90 °C and sonicated for 15 min. The coverslips were then rinsed with a copious amount of nanopure water, followed by incubation in a 1:1 (v/v) hydrochloric acid: methanol solution for 30 min at room temperature. After 30 min, the coverslips were rinsed multiple times with nanopure water and then dried overnight in a vacuum oven at 60 °C. The following day, the coverslips were cooled at room temperature and then incubated for 30 min in a 15% (v/v) solution of MPTMS in $N_2$ purged ethanol. The coverslips were sonicated for 10 min in $N_2$ purged ethanol three times and dried in a vacuum oven for at least 3 h at 120 °C. The coverslips were stored at 4°C for up to one week.

***Specific binding Hill slope equation***. Binding curves were fitted to a specific binding Hill slope to determine the $K_D$ values between $\alpha_2$ and PLN domains/subdomains. In this equation [BL] is the concentration of formed complexes between the binding sites (B, receptor) and the ligand (L, analyte). [B0] is the total concentration of binding sites, $K_D$ is the dissociation constant, $h$ is the Hill coefficient, and [L] is the concentration of the ligand[10-12].

$$\frac{[BL]}{[B_0]} = \frac{1}{1 + \frac{K_D}{[L]^h}}$$

### Supplementary Figures and Tables

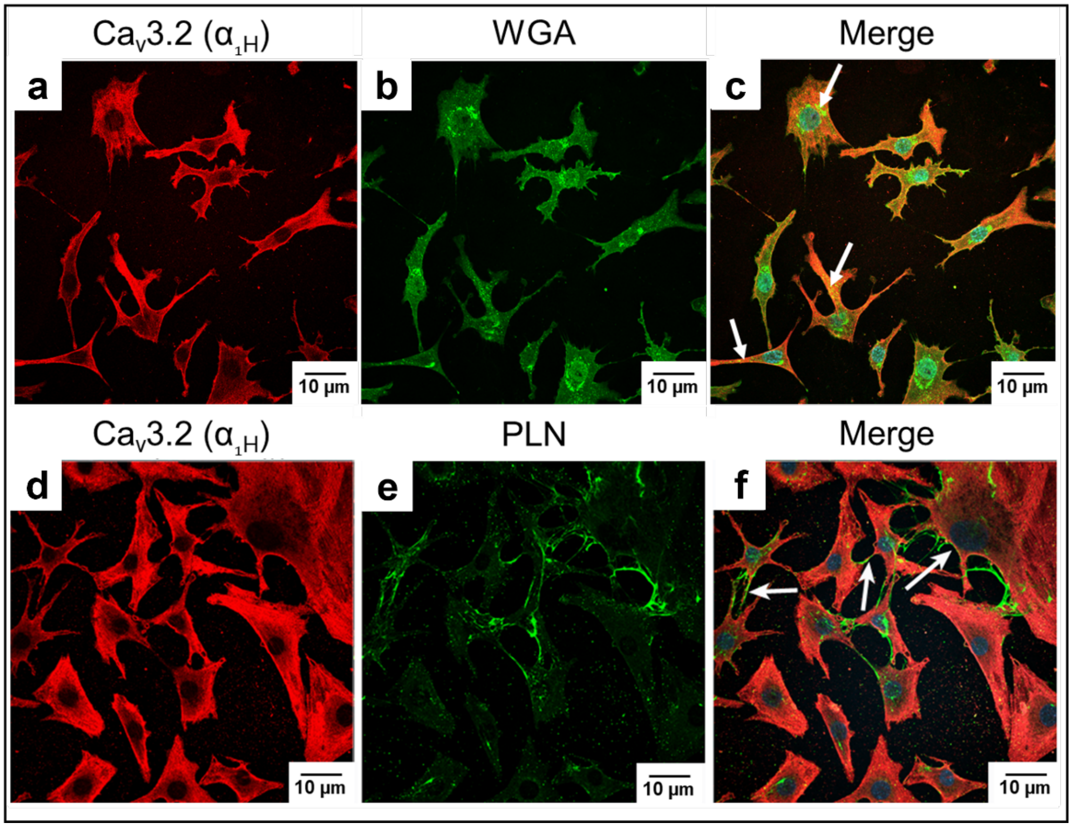


**Figure S1*. PLN and Ca$_v$3.2 ($\alpha_{1H}$) colocalize in osteocyte-like cells*.** Top. MLO-Y4 cells stained for **(a)** Ca$_v$3.2 ($\alpha_{1H}$) (red) and **(b)** fluorescein conjugated wheat germ agglutinin (WGA) (green). **(c)** The merged image shows overlapping fluorescent signal (white arrows) between $\alpha_{1H}$ and WGA. Bottom, Perlecan (PLN) and $\alpha_{1H}$ colocalize in osteocyte-like cells. MLO-Y4 cells stained for **(d)** Ca$_v$3.2 ($\alpha_{1H}$) (red) and **(e)** PLN (green). **(f)** The merged image shows overlapping fluorescent signal between $\alpha_{1H}$ and PLN (white arrows).

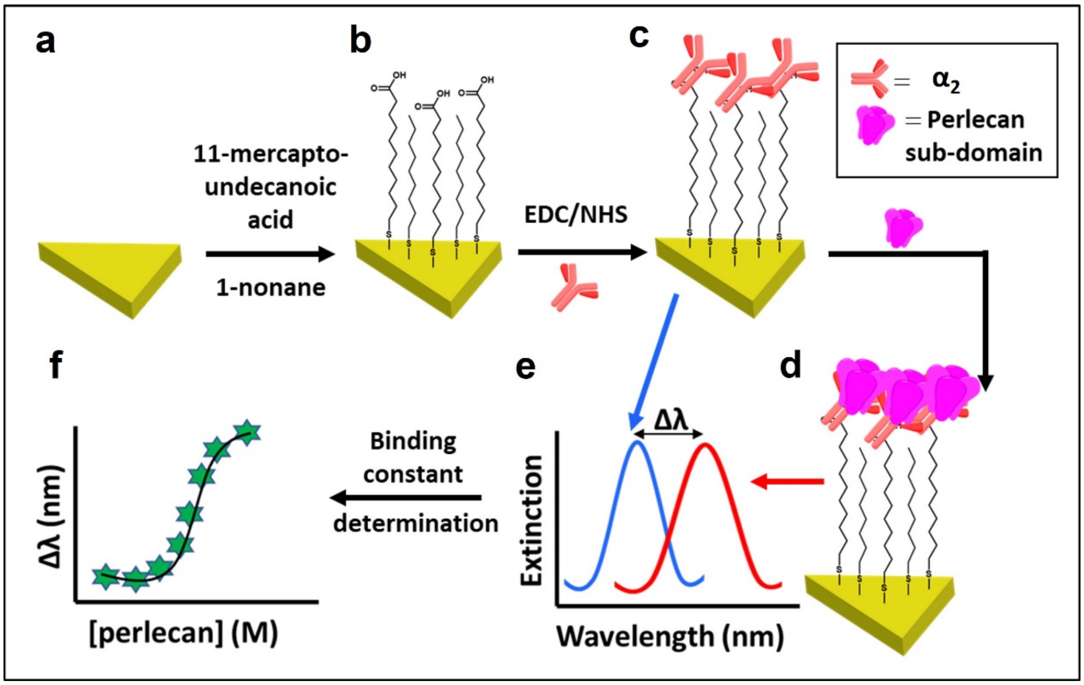


**Figure S2.** ***Schematic of $\alpha_2$-functionalized gold nanoprisms for LSPR binding affinity determination of PLN domains***. **(a)** Gold triangular nanoprisms were attached onto silanized glass coverslips. **(b)** Nanoprism surface was modified with mercaptoundecanoic acid and nonanethiol solution, and **(c)** the $\alpha_2$ portion of the $\alpha_2\delta_1$ subunit attached to the nanoprisms by amide coupling. **(d)** Perlecan (PLN) domains/subdomains were added, and **(e)** Localized surface plasmon resonance (LSPR) wavelengths were measured before (blue curve) and after incubation with PLN using UV-visible spectroscopy (red curve). **(f)** The plot of LSPR peak wavelength shift ($\Delta\lambda$) versus the log of PLN concentration (nM) was used to determine the binding affinity ($K_D$) by fitting the curve to Hill slope (dashed line).

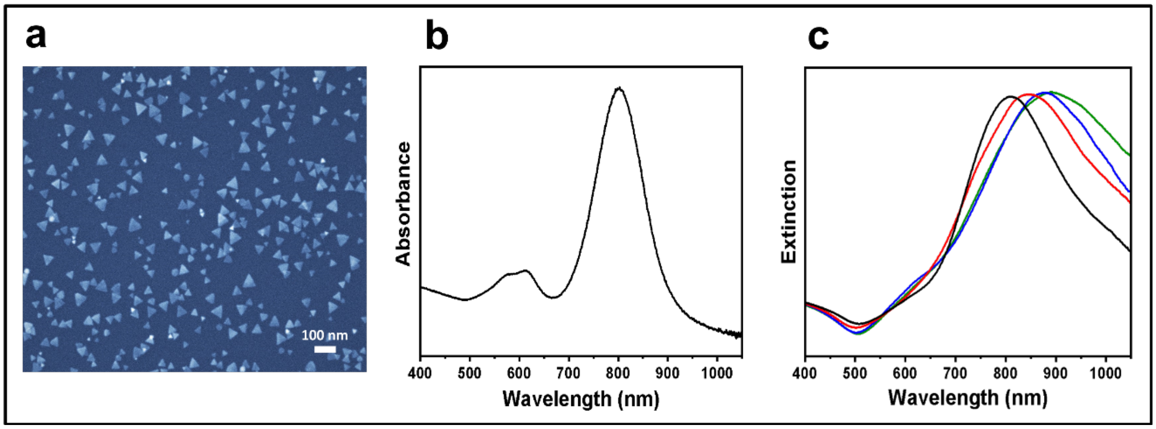


**Figure S3.** ***LSPR-based sensor characterizations*****.** **(a)** Scanning electron microscopy (SEM) image of ~42 nm edge length gold triangular nanoprisms (Au TNPs). **(b)** Representative UV-vis absorption spectrum of ~42 nm edge length Au TNPs in acetonitrile ($\lambda_{LSPR}$ = 801.9 nm). **(c)** UV-vis extinction spectra show an example of localized surface plasmon resonance (LSPR) shifts used to determine the binding affinity. UV−vis extinction spectra of silanized glass coverslip-bound Au TNPs before surface modification (black curve, 807.9 nm), after functionalization with a self-assembled monolayer (SAM) of 11-mercaptundecanoic acid (1.0 mM): 1-nonanethiol (1.0 μM) (red curve, 846.7 nm), after covalent attachment of 10 ng/mL $\alpha_2$ via amide coupling (blue curve, 885.7 nm), and finally after adsorption of perlecan domain III (green curve, 899.1 nm).

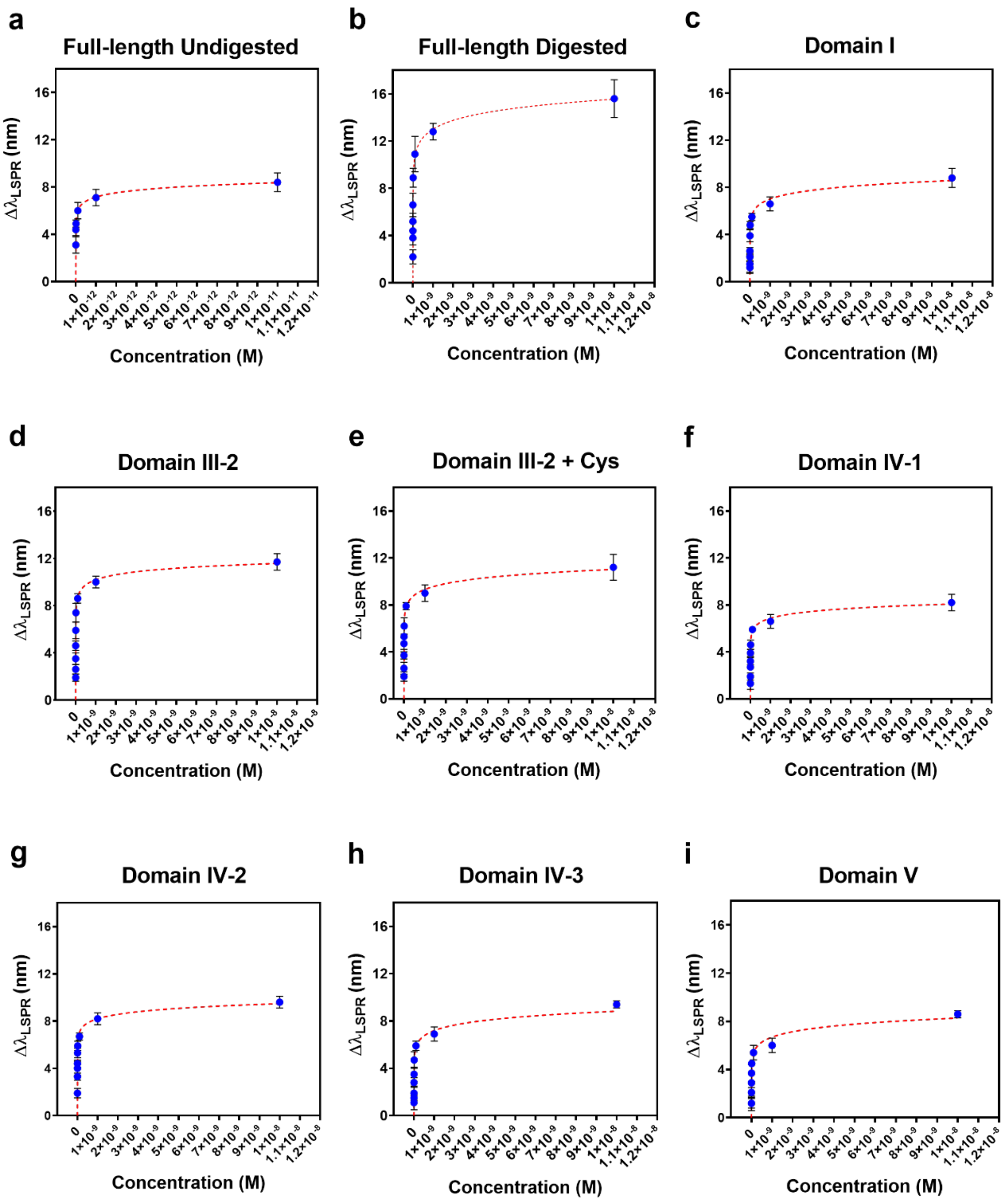


**Figure S4.** ***Binding Affinity between the*** $\boldsymbol{\alpha_2}$ ***portion of the*** $\boldsymbol{\alpha_2\delta_1}$ ***subunit and PLN***. $\alpha_2$-functionalized sensors were incubated with a solution containing different concentrations ($1 \times 10^{-16}$ to $1 \times 10^{-8}$ M) of full-length perlecan (PLN) or each of PLN domains (Dm). Dissociation constant curves were constructed for **(a)** Full-length PLN (undigested) and **(b)** enzymatically digested with heparinase and chondroitinase, **(c)** PLN Dm I **(d)** Dm III-2, **(e)** Dm III-2 with cysteine, **(f-h)** PLN Dm IV-1, -2, and -3, and **(i)** PLN Dm V. Each data point represents the average shift in the LSPR dipole peak position ($\Delta\lambda_{LSPR}$) values of six measurements (Mean± SD). Binding curves were developed by plotting $\Delta\lambda_{LSPR}$ versus PLN [or PLN subdomains] concentration in mol/L (M). Data from these curves was analyzed with the specific binding with Hill slope equation to determine $K_D$ values (8-10). The raw data corresponding to these graphs can be found in **Table S1**.

**Table S1.** *Raw data from binding affinity experiments between $\alpha_2\delta_1$ and Perlecan*

**a. PLN full length undigested**

| Concentration (M) | Full Length Undigested $\Delta\lambda_{LSPR}$ (nm) | Standard Deviation (nm) |
|---|---|---|
| $1x10^{-7}$ | - | - |
| $1x10^{-8}$ | - | - |
| $1x10^{-9}$ | - | - |
| $1x10^{-10}$ | - | - |
| $1x10^{-11}$ | 8.4 | 0.8 |
| $1x10^{-12}$ | 7.1 | 0.7 |
| $1x10^{-13}$ | 6 | 0.7 |
| $1x10^{-14}$ | 4.9 | 0.3 |
| $1x10^{-15}$ | 4.4 | 0.5 |
| $1x10^{-16}$ | 3.1 | 0.7 |

**b. PLN full length digested**

| Concentration (M) | Full Length Digested $\Delta\lambda_{LSPR}$ (nm) | Standard Deviation (nm) |
|---|---|---|
| $1x10^{-7}$ | - | - |
| $1x10^{-8}$ | 15.6 | 1.6 |
| $1x10^{-9}$ | 12.8 | 0.7 |
| $1x10^{-10}$ | 10.9 | 1.5 |
| $1x10^{-11}$ | 8.9 | 0.8 |
| $1x10^{-12}$ | 6.6 | 1 |
| $1x10^{-13}$ | 5.2 | 0.7 |
| $1x10^{-14}$ | 4.4 | 0.7 |
| $1x10^{-15}$ | 3.8 | 0.6 |
| $1x10^{-16}$ | 2.2 | 0.6 |

**c. PLN Dm I**

| Concentration (M) | Domain-I $\Delta\lambda_{LSPR}$ (nm) | Standard Deviation (nm) |
|---|---|---|
| $1x10^{-7}$ | - | - |
| $1x10^{-8}$ | 8.8 | 0.8 |
| $1x10^{-9}$ | 6.6 | 0.6 |
| $1x10^{-10}$ | 5.5 | 0.3 |
| $1x10^{-11}$ | 4.8 | 0.3 |
| $1x10^{-12}$ | 3.9 | 0.5 |
| $1x10^{-13}$ | 2.6 | 0.3 |
| $1x10^{-14}$ | 2.1 | 0.3 |
| $1x10^{-15}$ | 1.5 | 0.7 |
| $1x10^{-16}$ | 1.2 | 0.5 |

**d. PLN Dm III-2**

| Concentration (M) | Domain-III-2 $\Delta\lambda_{LSPR}$ (nm) | Standard Deviation (nm) |
|---|---|---|
| $1x10^{-7}$ | 13.4 | 1 |
| $1x10^{-8}$ | 11.7 | 0.7 |
| $1x10^{-9}$ | 10 | 0.5 |
| $1x10^{-10}$ | 8.6 | 0.4 |
| $1x10^{-11}$ | 7.4 | 0.8 |
| $1x10^{-12}$ | 5.9 | 0.7 |
| $1x10^{-13}$ | 4.6 | 0.4 |
| $1x10^{-14}$ | 3.5 | 0.5 |
| $1x10^{-15}$ | 2.6 | 0.4 |
| $1x10^{-16}$ | 1.9 | 0.3 |

**e. PLN Dm III-2 + Cysteine**

| Concentration (M) | Domain III-2 + cys $\Delta\lambda_{LSPR}$ (nm) | Standard Deviation (nm) |
|---|---|---|
| $1x10^{-7}$ | - | - |
| $1x10^{-8}$ | 11.2 | 1.1 |
| $1x10^{-9}$ | 9 | 0.7 |
| $1x10^{-10}$ | 7.9 | 0.3 |
| $1x10^{-11}$ | 6.2 | 0.7 |
| $1x10^{-12}$ | 5.3 | 0.2 |
| $1x10^{-13}$ | 4.7 | 0.5 |
| $1x10^{-14}$ | 3.7 | 0.3 |
| $1x10^{-15}$ | 2.6 | 0.5 |
| $1x10^{-16}$ | 1.9 | 0.4 |

**f. PLN Dm IV-1**

| Concentration (M) | Domain IV-1 $\Delta\lambda_{LSPR}$ (nm) | Standard Deviation (nm) |
|---|---|---|
| $1x10^{-7}$ | - | - |
| $1x10^{-8}$ | 8.2 | 0.7 |
| $1x10^{-9}$ | 6.6 | 0.6 |
| $1x10^{-10}$ | 5.9 | 0.3 |
| $1x10^{-11}$ | 4.6 | 0.4 |
| $1x10^{-12}$ | 3.9 | 0.3 |
| $1x10^{-13}$ | 3.2 | 0.3 |
| $1x10^{-14}$ | 2.7 | 0.2 |
| $1x10^{-15}$ | 1.9 | 0.3 |
| $1x10^{-16}$ | 1.3 | 0.5 |

**g. PLN Dm IV-2**

| Concentration (M) | Domain IV-2 $\Delta\lambda_{LSPR}$ (nm) | Standard Deviation (nm) |
|---|---|---|
| $1x10^{-7}$ | - | - |
| $1x10^{-8}$ | 9.6 | 0.5 |
| $1x10^{-9}$ | 8.2 | 0.5 |
| $1x10^{-10}$ | 6.7 | 0.3 |
| $1x10^{-11}$ | 5.9 | 0.4 |
| $1x10^{-12}$ | 5.3 | 0.4 |
| $1x10^{-13}$ | 4.4 | 0.3 |
| $1x10^{-14}$ | 4 | 0.6 |
| $1x10^{-15}$ | 3.3 | 0.3 |
| $1x10^{-16}$ | 1.9 | 0.4 |

**h. PLN Dm IV-3**

| Concentration (M) | Domain IV-3 $\Delta\lambda_{LSPR}$ (nm) | Standard Deviation (nm) |
|---|---|---|
| $1x10^{-7}$ | - | - |
| $1x10^{-8}$ | 9.4 | 0.3 |
| $1x10^{-9}$ | 6.9 | 0.6 |
| $1x10^{-10}$ | 5.9 | 0.4 |
| $1x10^{-11}$ | 4.7 | 0.7 |
| $1x10^{-12}$ | 3.5 | 0.6 |
| $1x10^{-13}$ | 2.8 | 0.4 |
| $1x10^{-14}$ | 1.9 | 0.6 |
| $1x10^{-15}$ | 1.5 | 0.4 |
| $1x10^{-16}$ | 1.1 | 0.6 |

**i. PLN Dm V**

| Concentration (M) | Domain-V $\Delta\lambda_{LSPR}$ (nm) | Standard Deviation (nm) |
|---|---|---|
| $1x10^{-7}$ | - | - |
| $1x10^{-8}$ | 8.6 | 0.3 |
| $1x10^{-9}$ | 6 | 0.6 |
| $1x10^{-10}$ | 5.4 | 0.6 |
| $1x10^{-11}$ | 4.5 | 0.2 |
| $1x10^{-12}$ | 3.7 | 0.7 |
| $1x10^{-13}$ | 2.9 | 0.2 |
| $1x10^{-14}$ | 2.1 | 0.4 |
| $1x10^{-15}$ | 1.2 | 0.4 |
| $1x10^{-16}$ | 1.2 | 0.6 |
